# Chromatin accessibility profiling uncovers genetic- and T2D disease state-associated changes in *cis*-regulatory element use in human islets

**DOI:** 10.1101/192922

**Authors:** Shubham Khetan, Romy Kursawe, Ahrim Youn, Nathan Lawlor, Eladio Marquez Campos, Duygu Ucar, Michael L. Stitzel

## Abstract

Genetic and environmental factors both contribute to islet dysfunction and failure, resulting in type 2 diabetes (T2D). The islet epigenome integrates these cues and can be remodeled by genetic and environmental variation. However, our knowledge of how genetic variants and T2D disease state alter human islet chromatin landscape and *cis-*regulatory element (RE) use is lacking. To fill this gap, we profiled and analyzed human islet chromatin accessibility maps from 19 genotyped individuals (5 with T2D) using ATAC-seq technology. Chromatin accessibility quantitative trait locus (caQTL) analyses identified 3001 sequence variants (FDR<10%) altering putative *cis-*RE use/activity. Islet caQTL were significantly and specifically enriched in islet stretch enhancers and islet-specific transcription factor binding motifs, such as FOXA2, NKX6.1, RFX5/6 and PDX1. Importantly, these analyses identified putative functional single nucleotide variants (SNVs) in 13 T2D-associated GWAS loci, including those previously associated with altered *ZMIZ1, MTNR1B, RNF6,* and *ADCY5* islet expression, and linked the risk alleles to increased (n=8) or decreased (n=5) islet chromatin accessibility. Luciferase reporter assays confirmed allelic differences in *cis-*RE activity for 5/9 caQTL sequences tested, including a T2D-associated SNV in the *IL20RA* locus. Comparison of T2D and non-diabetic islets revealed 1882 open chromatin sites exhibiting T2D-associated chromatin accessibility changes (FDR<10%). Together, this study provides new insights into genetic variant and T2D disease state effects on islet *cis-*RE use and serves as an important resource to identify putative functional variants in T2D-and islet dysfunction-associated GWAS loci and link their risk allele to *in vivo* loss or gain of chromatin accessibility.

## Introduction

Pancreatic islet function is central to maintaining glucose homeostasis. T2D is a complex disease resulting from the combined effects of genetic susceptibility and environmental exposures. Genome-wide association studies (GWAS) have associated single nucleotide variants (SNVs) in >100 loci with increased susceptibility to type 2 diabetes (T2D) and related quantitative measures of islet dysfunction (Fuchsberger et al. 2016; Mohlke and Boehnke 2015). The majority of these variants overlap islet-specific enhancer elements (Parker et al. 2013; Fuchsberger et al. 2016; Pasquali et al. 2014). This enrichment establishes perturbed islet transcriptional regulation in the genetic etiology of islet dysfunction and T2D (Lawlor et al. 2017). In addition to individual genetic variation, environmental insults to islet functions such as oxidative, endoplasmic reticulum (ER), and inflammatory stresses, have also been linked to T2D.

Genetic and environmental factors shape the epigenome to modulate the transcriptional programs governing steady state and stress responsive-factors. Common genetic variants in the human population, contributing to complex phenotypes and disease susceptibility, have been linked to alterations in regulatory element use, as monitored by changes in chromatin accessibility (Degner et al. 2012; Pique-Regi et al. 2011; Kumasaka et al. 2016; McDaniell et al. 2010; Alasoo et al. 2017) and histone modifications (McVicker et al. 2013; Ng et al. 2017) in diverse cell types. Moreover, *cis*-regulatory element use in a given cell type can be modified by its local environment (Lavin et al. 2014) and cellular responses to stimuli (Ostuni et al. 2013) and stressors (Brown et al. 2014). Currently, our understanding of how individual genetic variation and the type 2 diabetic disease state alter *cis*-regulatory element use in human pancreatic islets is limited.

In this study, we sought to understand how (1) genetic variants, particularly those associated with T2D susceptibility and quantitative measures of islet dysfunction; and (2) T2D state alter chromatin accessibility and *cis*-regulatory element use in human islets. Using the assay for transposase-accessible chromatin with sequencing (ATAC-seq), we studied the chromatin accessibility patterns in human islets obtained from 19 individuals, five of whom were type 2 diabetic. Using RASQUAL (Kumasaka et al. 2016), we integrated genotypes generated in each individual with their corresponding open chromatin profiles to identify chromatin accessibility quantitative trait loci (caQTLs), i.e., genetic variants that alter chromatin accessibility and *cis*-RE use in islets. Finally, by comparing ATAC-seq profiles between diabetic and normal donors, we identified changes in chromatin accessibility and putative *cis*-RE use associated with the T2D disease state.

## Results

### Chromatin accessibility maps in human pancreatic islets

To determine the genome-wide location of *cis*-regulatory elements in human islets, we transposed the nuclei of islet samples obtained from 23 cadaveric organ donors (Table 1; 18 non-diabetic (ND) and 5 T2D) and measured chromatin accessibility using ATAC-seq. 19/23 (n=14 ND, 5 T2D) donor islet ATAC-seq datasets passed quality control filters (Methods) and were used in subsequent analyses (Figures 1A, S1A; Supplemental Table 1). As shown in Figure 1B, the genome-wide chromatin accessibility profiles of these islets were highly correlated and all islet profiles clustered separately from other cell types, including T2D-relevant skeletal muscle and adipose tissues. Notably, T2D donor islet profiles (n=5) did not cluster distinctly from those of ND donor islets (n=14), suggesting that the T2D disease state itself does not lead to global restructuring of islet chromatin accessibility. ATAC-seq profiles of representative islets for the *GCK* locus are shown in Figure 1C, revealing both common (gray) and islet-specific (orange) ATAC-seq peaks.

**Figure 1.**
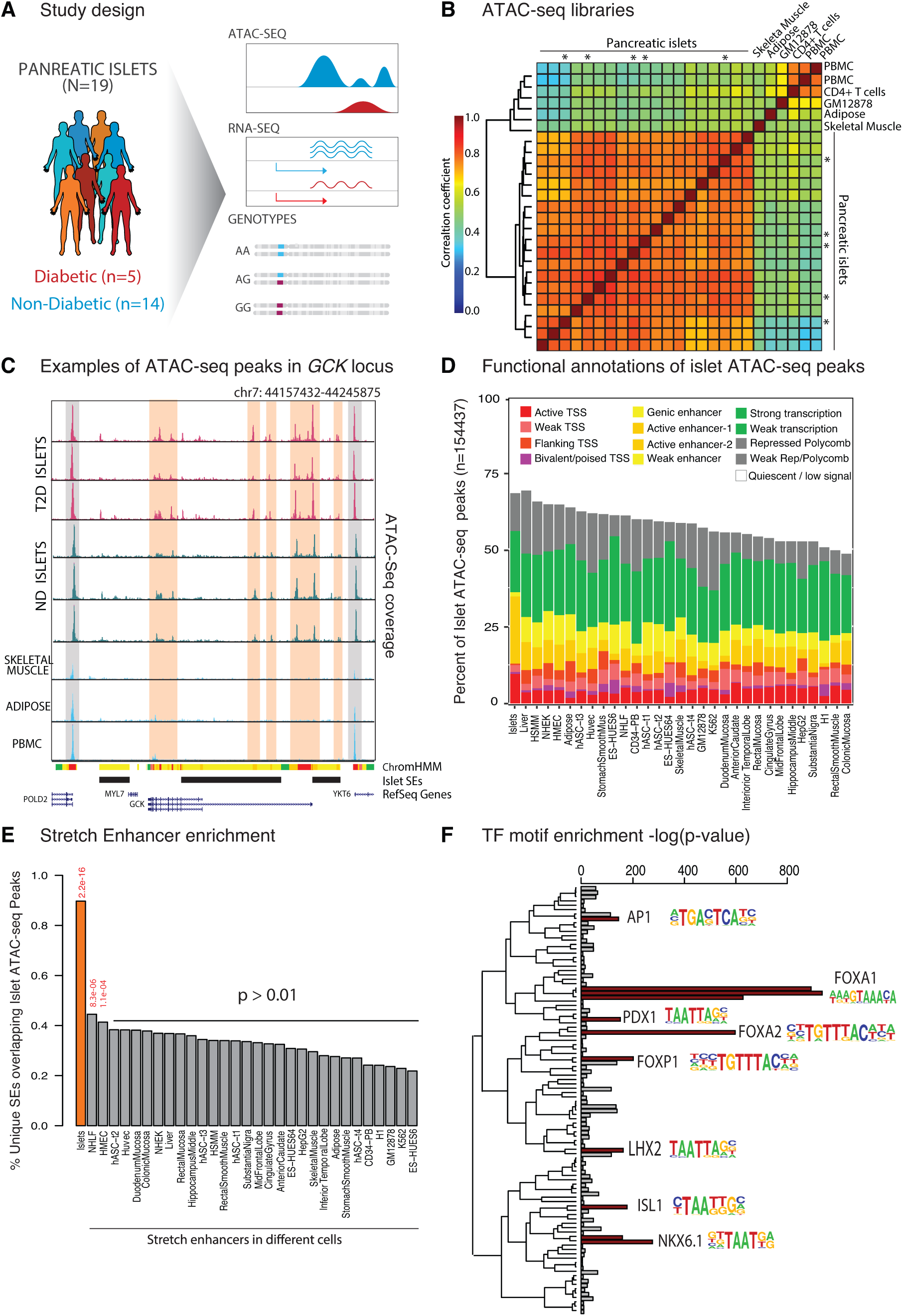
Chromatin accessibility maps in human islets. **(A)** Schematic of study design. **(B)** Spearman correlation between genome-wide ATAC-seq read distributions of islets and other cell types. Islets from type 2 diabetic donors are marked with asterisk. (PBMC->peripheral blood mononuclear cells). **(C)** An example locus in and around the GCK gene representing chromatin accessibility landscapes in 3 ND and 3 T2D islets, and other tissues. The regions marked in orange are specifically accessible in islet cells, whereas regions marked in gray are ubiquitously accessible. All chromatin accessibility maps are normalized to the same depth and have the same scale. **(D)** Overlap of islet ATAC-seq peaks with chromatin states in islets and 30 other cell types. Tissues are sorted from highest to lowest overlap between ATAC-seq peaks and enhancer states.TSS = Transcription Start Site). **(E)** Percent of tissue-specific stretch enhancers (SEs) overlapping islet ATAC-seq peaks. Fisher’s exact test p-values are shown to represent enrichment. **(F)** Enrichment of transcription factor (TF) motifs in islet-specific ATAC-seq peaks.TFs are clustered with respect to the similarity of their position weight matrices (PWMs).

Collectively, 154,438 ATAC-seq peaks were identified across the 19 islet donors (see Methods), representing putative *cis*-REs (i.e., promoter, enhancer, repressor, and insulator elements). 40% of these putative islet REs were detected in only 1 out of 19 individuals in the cohort (Supplementary Figure S1B). Not surprisingly, 45% of these individual-specific peaks were un-annotated in reference islet ChromHMM states (i.e., low signal state) (Supplementary Figure S1C). In contrast, ATAC-seq peaks at gene promoters were consistently accessible across the cohort (Supplementary Figure S1C), suggesting that promoter elements are less variable across individuals compared to other *cis*-REs.

Islet ATAC-seq peaks were compared against previously reported (Varshney et al. 2017) ChromHMM-defined functional states in islets from our previous studies (Stitzel et al. 2010; Parker et al. 2013) and 30 other tissues from the NIH Epigenome Roadmap project (Roadmap Epigenomics Consortium et al. 2015), including adipose, skeletal muscle, liver, and brain. As anticipated, islet ATAC-seq peaks were most enriched in ChromHMM-defined islet enhancers (Figure 1D). Similarly, we compared the islet ATAC-seq peaks with stretch enhancers (SEs) in 31 tissues. SEs are long (>3kb) stretches of contiguous cell-specific enhancer chromatin states that are enriched for disease-associated SNVs relevant to the cognate cell type (Parker et al. 2013). 90% of islet SEs overlapped islet ATAC-seq peaks (Figure 1E); this overlap was significantly greater (Fisher’s Exact p<2.2e-16 for islet SEs) than that observed for other tissue SEs. Moreover, islet-specific peaks (i.e., those that do not overlap open chromatin sites in Skeletal Muscle, Adipose, GM12878, CD4+ T cells, PBMCs), were enriched in motifs of islet cell-specific transcription factors, such as PDX1 and NKX6.1, when compared to ATAC-seq peaks that were common across tissues (Figure 1F). These data thus represent high quality chromatin accessibility maps of human islets and captures islet-specific regulatory elements.

### Identification of genetic variants affecting islet chromatin accessibility

Since genetic variation in *cis*-RE use/activity has been implicated in diverse phenotypes and complex diseases, including T2D, we sought to identify genetic variants that alter chromatin accessibility in human islets (Figure 2A). Chromatin accessibility QTL (caQTL) analysis (Kumasaka et al. 2016) was used to identify the genetic variants within each ATAC-seq peak that correlated with changes in its accessibility (see Methods). In total, we uncovered 3001 SNVs significantly associated (FDR<0.10; Fig S2A) with chromatin accessibility changes in this cohort. Figure 2B highlights an example caQTL overlapping an intronic islet SE in the *CELF4* gene, which exhibits islet-selective expression (Varshney et al. 2017). Islet chromatin accessibility was reduced in rs488797 CC homozygotes, potentially by disrupting a FOXA2 binding motif (Ward and Kellis 2016). In agreement, almost all ATAC-seq reads overlapping this variant in CT heterozygous islets contain the T allele (Figure 2B, inset).

**Figure 2.**
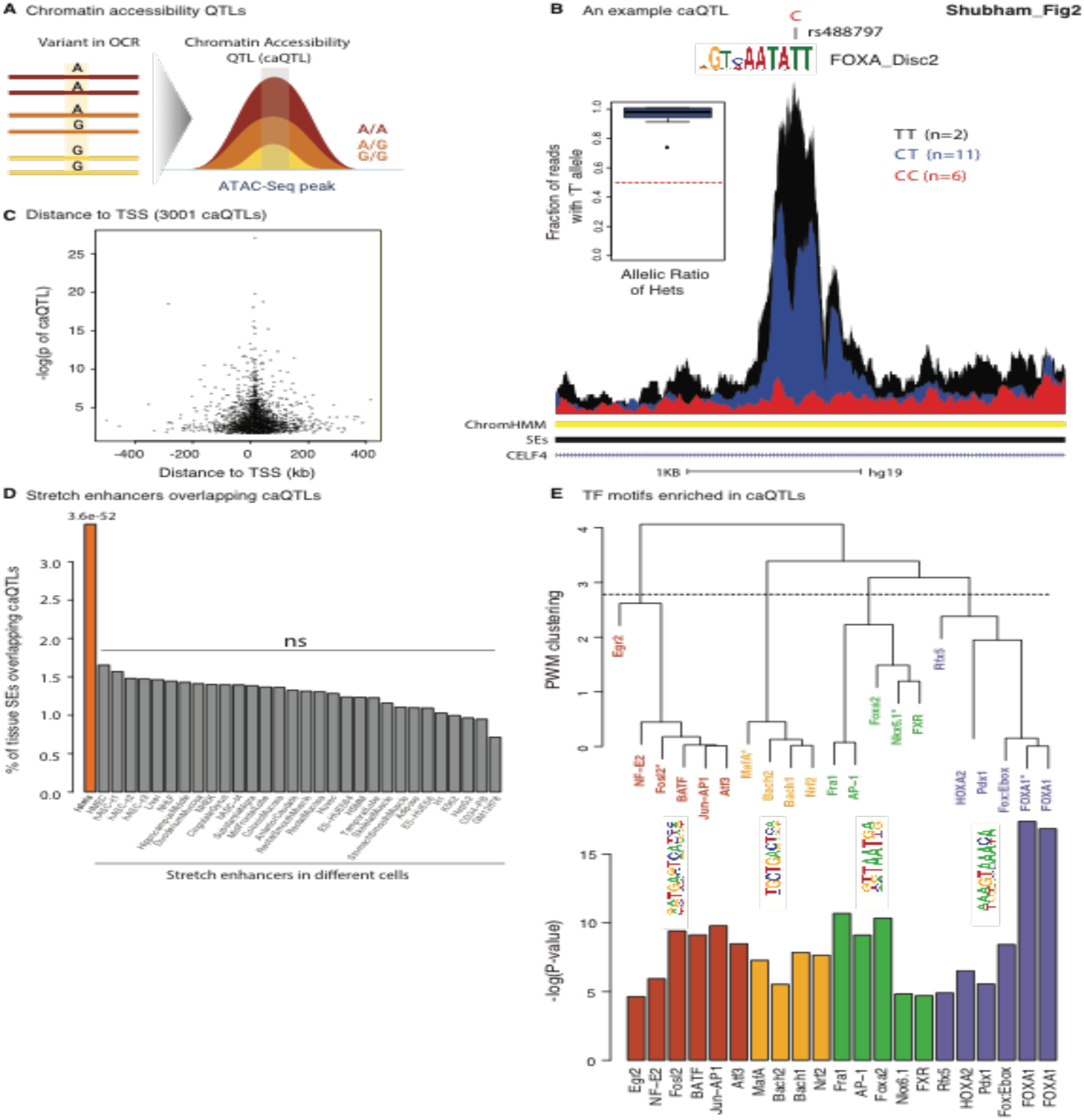
Identification of genetic variants affecting islet chromatin accessibility. **(A)** Schematic of chromatin accessibility quantitative trait locus (caQTL) analyses linking genotype to chromatin accessibility changes. (**B**) An example caQTL in an islet-SE within an intron of *CELF4*. Aggregate ATAC-seq profiles of individuals with C/C (red), C/T (blue), and T/T (black) genotypes at rs488797 are displayed. The inset boxplot shows the fraction of ATAC-seq reads containing the T allele at rs488797 in each C/T heterozygous islet (n=11). Sequence logo for FOXA2 TF is displayed, which is predicted by HaploReg to be altered by this variant. (**C**) Distance between caQTLs and the transcription start site (TSS) of the nearest expressed gene versus the significance of the caQTL association. The majority of caQTLs are within 200 kb of the TSS of the nearest expressed gene. (**D**) Percent of tissue-specific SEs overlapping caQTLs. Fisher’s exact test p-values are shown for enrichment. (**E**) TF motifs enriched in islet caQTLs. TFs are clustered based on their PWM similarity using hierarchical clustering, resulting in four major TF groups. Bar plots of p-values are color coded according to this clustering. A representative PWM logo is represented for each cluster, where the corresponding TF is marked with an asterisk.

The majority of islet caQTLs (97%) were within 200 kilobases (kb) of the transcription start site (TSS) of an islet-expressed gene. (Figure 2C). Approximately 30% of islet caQTLs overlapped islet enhancer chromatin states (Supplementary Figure S2C). When compared to tissue SEs, the islet caQTLs were specifically enriched in islet SEs only (Figure 2E). Peaks containing caQTLs were also significantly enriched in islet SEs when compared to all islet ATAC-seq peaks (p=0.0024; OR=1.18; Fisher’s exact test), suggesting that islet caQTLs alter *cis*-REs encoding important islet-specific functions. Consistently, sequence motifs for islet-specific TFs, such as NKX6.1, PDX1, and MAFA, and not for general TFs, were enriched in caQTL-containing ATAC-seq peaks (Figure 2F). Surprisingly, sequence motifs of oxidative stress-responsive TFs (Ma 2013; Dhakshinamoorthy et al. 2005), such as BACH1, BACH2, and NRF2, were also enriched in caQTL peaks. Together, these data and analyses enumerate sequence variants that alter chromatin accessibility of islet *cis*-REs and suggest that these changes may be associated with altered binding of TFs governing islet cell identity, function, and stress response.

### T2D-associated GWAS SNVs alter chromatin accessibility in islets

The large majority (>90%) of common variants associated with T2D and quantitative measures of islet dysfunction, such as fasting plasma glucose and insulin levels, reside in non-coding loci and significantly and specifically overlap islet SEs. However, *in vivo* effects of these GWAS SNV alleles on *cis*-regulatory element use, as assessed by chromatin accessibility, islet TF ChIP, or active enhancer histone modifications such as H3K27ac, have been assessed and reported at only a handful of loci (Gaulton et al. 2010; Roman et al. 2017) to date. We hypothesized that T2D-and islet (dys)function-associated GWAS SNVs alter chromatin accessibility in islets, and exhibit significant and specific overlap with islet caQTLs. To test this, we assessed the enrichment (Schmidt et al. 2015) of islet caQTLs for SNVs exhibiting genome-wide significant (p<5e^−8^) associations with 184 diverse traits and diseases retrieved from the NHGRI/EBI GWAS Catalog (Methods). Among all GWAS traits and diseases assessed, islet caQTLs only exhibited significant enrichment of GWAS SNVs associated with T2D (2.97-fold), fasting glucose (13.46-fold), and BMI-adjusted fasting glucose (7.43-fold) (Figure 3A, p<5.43e-04, < 0.10 after Bonferroni correction).

**Figure 3.**
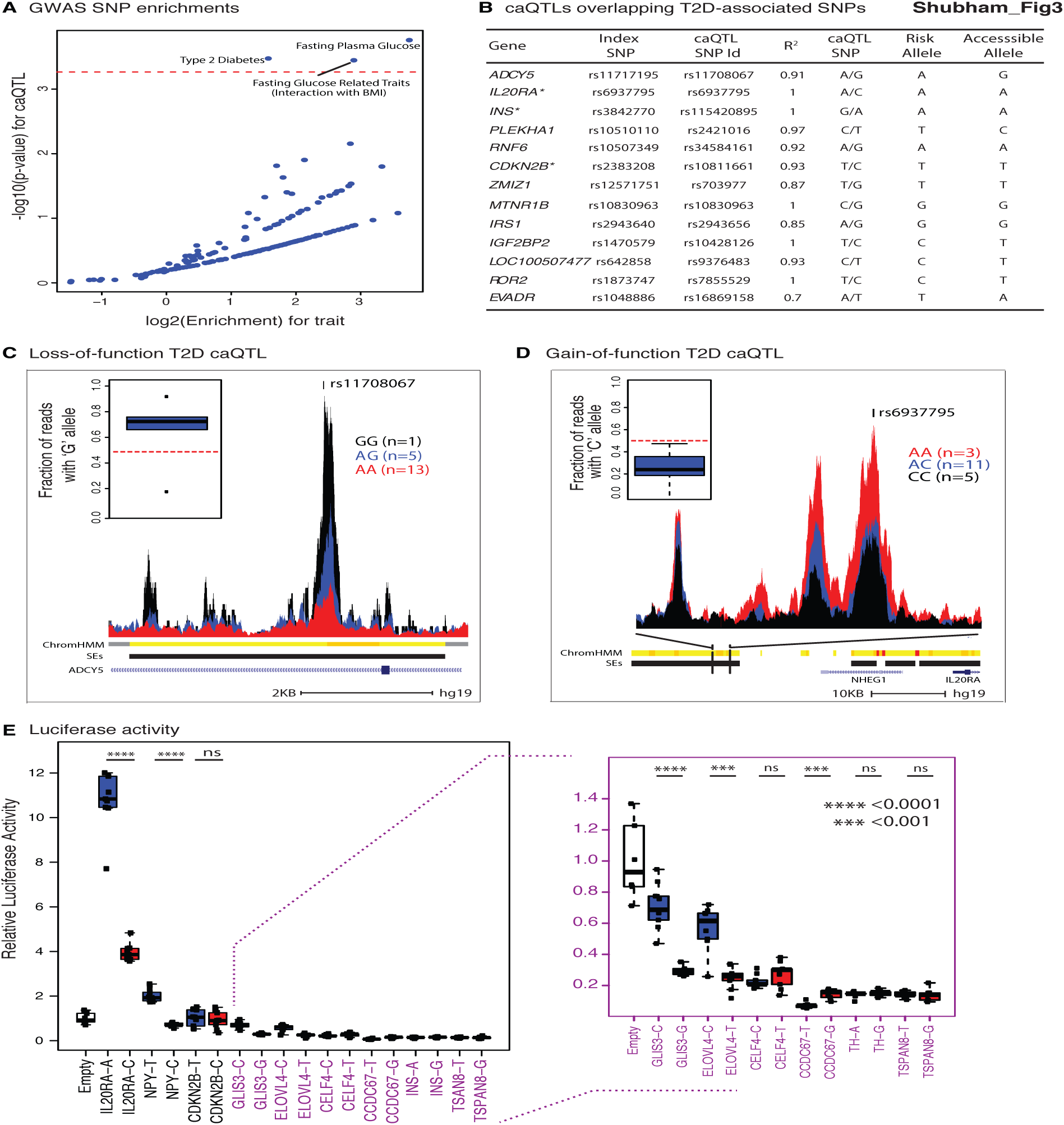
T2D associated caQTLs. **(A)** Enrichment of caQTLs for disease associated GWAS SNVs. The dashed red line represents p=5.43e^−4^, equivalent to 10% cutoff after Bonferroni correction for the number of diseases tested (n=184). (**B**) Table enumerating the 13 caQTLs linked to T2D-associated GWAS SNVs. Asterisks mark those loci that were tested for differential luciferase activity in panel E. (**C**) Average chromatin accessibility profiles at the *ADCY5* locus (loss-of-function T2D caQTL). The inset boxplot shows the fraction of ATAC-seq reads containing the G allele in each of the heterozygous islet samples (n=5). (**D**) Average chromatin accessibility profiles at the *IL20RA* locus (gain-of-function T2D caQTL). The inset boxplot shows the fraction of ATAC-seq reads containing the C allele in each of the heterozygous islet samples (n=11). (**E**) Luciferase activity of 9 tested caQTLs with reference and alternate alleles (normalized to empty construct) ^****^ and ^***^ indicate p<0.0001 and p<0.001, respectively; two-sided Mann-Whitney test p-values are shown on boxplots. ns = not sinificant.

These analyses highlighted 13 T2D-associated variants overlapping islet caQTLs (Figure 3B). They included 4 loci (*ADCY5, ZMIZ1, MTNR1B,* and *RNF6*) in which the caQTL SNV has been previously linked to altered *in vitro* enhancer activity or *in vivo* steady state gene expression in islets (van de Bunt et al. 2015; Roman et al. 2017; Lyssenko et al. 2009, 1; Varshney et al. 2017; Fadista et al. 2014). Importantly, for all loci that harbor both islet caQTL and eQTL variants (i.e., *ADCY5, ZMIZ1, MTNR1B,* and *RNF6*), the risk alleles have a concordant effect both on chromatin accessibility and gene expression levels (Figure 3B).

For 5 out of these 13 GWAS loci, the risk allele decreased chromatin accessibility (Figure 3B). This includes the T2D-associated index SNV rs11708067, which resides in the third intron of the *ADCY5* gene and overlaps an islet SE. The risk allele for this variant (A) is associated with reduced chromatin accessibility (Figures 3B, 3C). This is consistent with recent reports by us and others linking the rs11708067 risk allele (A) to decreased enhancer activity in luciferase reporter assays *in vitro* (Roman et al. 2017), to reduced histone H3 lysine 27 acetylation (H3K27ac) (Roman et al. 2017) and to decreased *ADCY5* expression in human islets *in vivo* (van de Bunt et al. 2015; Varshney et al. 2017; Roman et al. 2017). In the remaining 8 T2D-associated caQTL loci (Figures 3B, D), the T2D risk allele was associated with higher chromatin accessibility than the non-risk allele, suggesting that the risk allele is associated with a gain-of-function.

To validate a subset (n=9) of the islet caQTLs, we tested whether the human caQTL alleles altered enhancer activity of the sequences overlapping these putative *cis*-REs using luciferase reporter assays in MIN6 beta cells. Comparison of sequences containing either the reference or alternate allele for each caQTL site (Table 4) confirmed differential enhancer activity for 5 out of 9 loci (Figure 3E). For example, the rs6937795 “A” allele in the *IL20RA* locus, which is associated with increased T2D susceptibility and increased islet chromatin accessibility (Figure 3D), showed 2.5-fold higher enhancer activity than the non-risk “C” allele (Figures 3B, 3D, 3E).

### Chromatin accessibility changes in T2D versus ND islets

To uncover T2D associated changes in chromatin accessibility, we compared chromatin accessibility maps from 5 ND and 5 T2D donors (Figure 4A). Out of 52,387 ATAC-seq peaks tested, 1882 differentially accessible peaks between T2D and ND islets were identified (FDR 10%). Of these, 980 showed an increase and 902 showed a decrease in accessibility with the T2D state, hereafter referred to as “opening” and “closing” peaks, respectively (Figure 4B). There was a remarkable difference in the functional annotation of differential peaks, where closing peaks were mostly found at enhancers (48%), and opening peaks were mainly at promoters (70%) (Figure 4C). Figure 4D and 4E represents examples of closing and opening peaks respectively.

**Figure 4.**
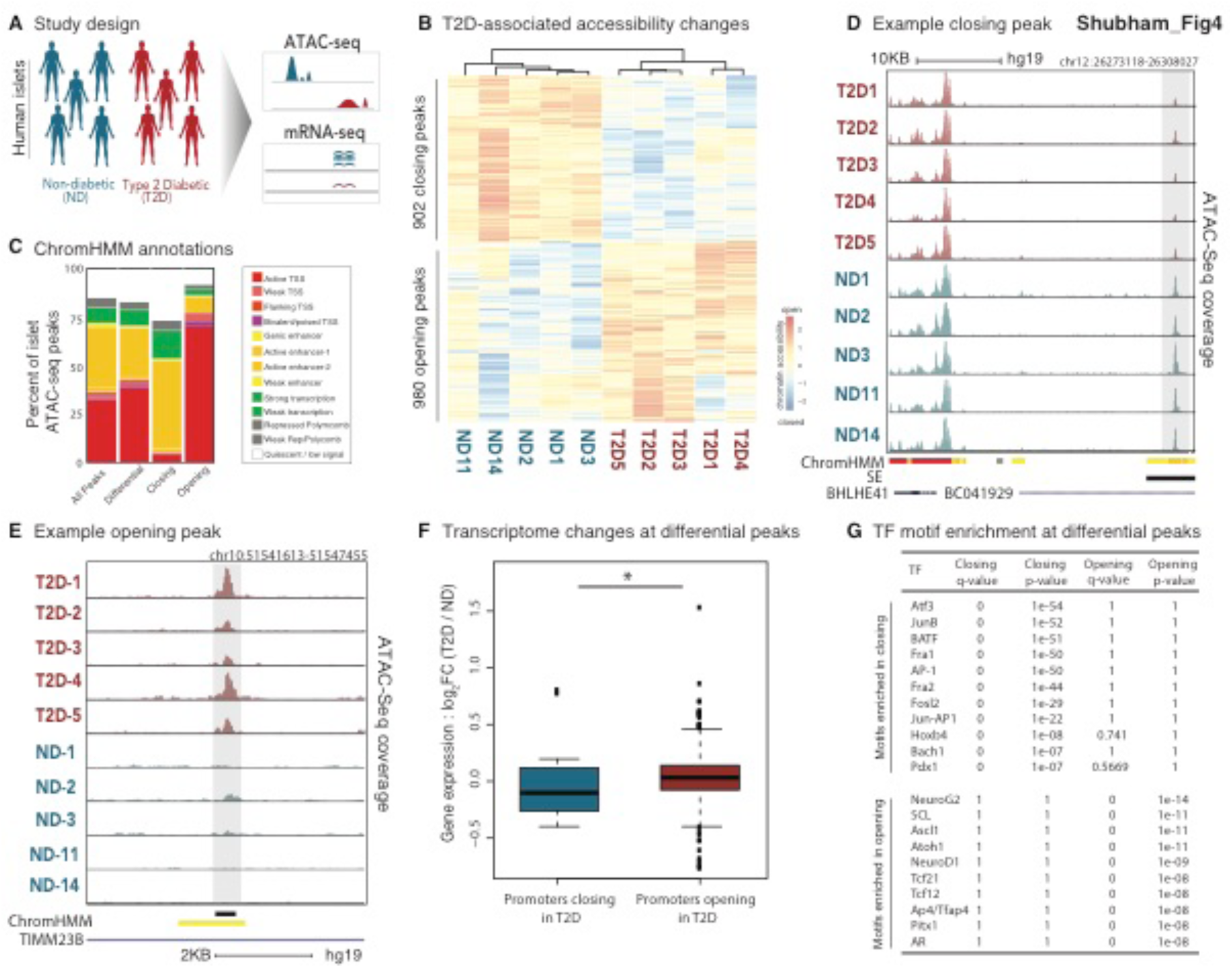
T2D-associated chromatin accessibility changes. **(A)** Schema summarizing the comparison of 5 T2D and 5 ND islets for chromatin accessibility and gene expression changes. (**B**) T2D-associated chromatin accessibility changes at differentially accessible sites (FDR 10%). Heat map represents normalized chromatin accessibility levels. (**C**) Islet ChromHMM annotations for ATAC-seq peaks (n=52387), differential peaks (n=1882), opening (n=980) and closing (n=902) peaks. Note that closing peaks are primarily at enhancer elements, whereas opening peaks are at promoter peaks. (**D**) Chromatin accessibility levels for an example loci around the *BHLHE41* gene that contains a closing peak (marked in gray bar). (**E**) Chromatin accessibility levels for an example loci in the *TIMM23B* gene that contains an opening peak. (**F**) Gene expression changes (measured in log fold change) for promoter peaks that are closing or opening in T2D islets. Note that genes with opening peaks have positive fold change, i.e., increased expression with T2D, whereas genes with closing peaks have negative fold change, i.e., decreased expression in T2D (p=0.038; Mann-Whitney test). (**G**) TF motif enrichment p-values for opening and closing peaks.

However, when differential peaks were categorized with respect to the presence or absence of ATAC-seq peaks in normal and diabetic donors (Fig S3E), we found that the majority of T2D-associated changes in chromatin accessibility were gradient in nature, i.e., peaks do not completely appear/disappear in T2D islets, with a few exceptions (<1%) (Supplementary Figure S3E). Additionally, we note that a subset of the 1882 differentially accessible peaks (42 opening peaks, 51 closing peaks) overlapped caQTLs (Supplementary Figure S3F), suggesting that these T2D associated accessibility changes are driven by genetic factors.

The differential peaks were annotated to the closest active genes in islets (Methods), resulting in 898 genes associated with opening and 665 genes associated with closing peaks (Supplemental Table S5). Differential gene expression analyses between ND and T2D samples revealed small changes in gene expression levels, where only 90 (38) genes were significantly up (down) regulated in T2D islets (FDR 10%). We observed a modest yet positive correlation between T2D-associated chromatin accessibility changes at gene promoters and the changes in the expression levels of these genes (p=0.038, Wilcoxon, Figure 4F). Opening and closing peaks were enriched in different TF motifs (Figure 4G). Interestingly, TFs that regulate stress responses such as ATF3, AP-1 were enriched in closing peaks.

## Discussion

Using ATAC-seq, we profiled the chromatin accessibility of human islets from 19 individuals. Integration of these open chromatin maps with each individual’s genotypes identified 3001 sequence variants (caQTL) that modulate *in vivo* islet regulatory element use. These caQTLs were enriched in islet-specific enhancers and TF motifs. Importantly, a subset of these was significantly and specifically enriched for T2D and fasting glucose GWAS index or linked (r^2^>0.8) sequence variants. Comparison of ATAC-seq profiles from T2D and ND individuals revealed quantitative changes in chromatin accessibility at 1882 putative islet *cis*-REs. Together, these data and analyses contribute significantly to (1) enumerating the genetic variants that alter islet *cis*-RE use; (2) delineating putative functional variants among the T2D- and islet dysfunction-associated GWAS SNVs; (3) linking the risk allele to *in vivo* loss or gain of *cis*-RE use in islets; and (4) assessing the relative chromatin accessibility effects of genetic variation and T2D state on human pancreatic islets.

ATAC-seq profiling in islets obtained from multiple cadaveric organ donors identified genetic effects on accessibility of 3.5% (3001/84499) of putative islet *cis*-REs genome-wide, and linked the alternate allele to increased chromatin accessibility at 43.5% (1307/3001) of sites and decreased accessibility in 56.5% (n=1694/3001) of sites compared to the reference allele. 6.2% (187/3001) of caQTLs variants identified in this study were in linkage disequilibrium (r^2^>0.8) with previously described islet eQTL (Varshney et al. 2017). Reports studying other cell types from larger cohorts have observed overlaps between chromatin-based QTLs (such as DNase-sensitivity and histone acetylation) and eQTLs ranging from 16%-45% (Degner et al. 2012; Li et al. 2016; del Rosario et al. 2015; Ng et al. 2017). Lower overlap observed in our study could be explained, at least in part, by the differences in sample size and resulting disparities in power between these two studies. It may also reflect the effect of the mixed cellular composition of islets, which might be resolved by studies measuring these features in sorted cell types. Finally, this overlap may reflect different inherent features measured by RNA-seq and chromatin-based assays that may contribute to these modest overlaps. For example, features reflected in RNA-seq data such as mRNA stability, polyadenylation, and splicing are not captured by chromatin profiling assays. This warrants future studies examining the impact of genetic variation on the ability of islets to respond to environmental changes (response QTLs), alongside islet caQTL studies with higher sample sizes, perhaps in sorted cells.

Using luciferase assays, we assessed allelic effects of the sequences overlapping nine of the islet caQTL sites on enhancer activity. Only 3/9 sequences tested exhibited transcriptional enhancer activity compared to the minimal promoter sequence alone, reinforcing the concept that caQTLs capture both enhancer and repressor *cis*-REs (Petrykowska et al. 2008). Importantly, five of these sites exhibited significant allelic differences in *cis*-RE activity. In each case, the direction of allelic effect on enhancer activity matched the allelic changes in chromatin accessibility, including for the rs6937795 variant in the T2D-associated *IL20RA* locus. For the remaining four sequences, lack of allelic differences in *in vitro* enhancer activity may be due to the human enhancers not being active in the mouse MIN6 beta cell line used for luciferase assays or to REs displaying enhancer activity only under certain conditions, such as oxidative stress and not in baseline conditions. Indeed, studies in other cell types suggest that regulatory elements can be primed for and activated by specific environmental stimuli or stressors (Ostuni et al. 2013; Alasoo et al. 2017; Brown et al. 2014).

In this study, we identified SNVs in 13 T2D-associated loci that alter chromatin accessibility. These include four loci (*ZMIZ1, MTNR1B, RNF6,* and *ADCY5*) in which the same or linked (r2>0.8) genetic variant has been identified as an islet eQTL. Importantly, the caQTL and eQTL studies identified a consistent direction-of-effect (e.g., gain-or loss-of-function) for the risk allele in each of these loci. T2D risk alleles in 5/13 loci were associated with reduced chromatin accessibility. For the remaining loci, the risk alleles were associated with increased chromatin accessibility, representing potential gain-of-function variants. Unfortunately, sequence motif analysis of these caQTL variants did not reveal obvious *trans*-factors that may be responsible for these accessibility differences or whose binding is affected by this sequence variant. This could be due in part to incomplete information on position weight matrices for TFs, including ARX, which is an islet alpha cell transcription factor. Together, these data and analyses have identified novel SNV effects on islet *cis*-REs, including their direction-of-effect, that can be further dissected in a site-specific and hypothesis-driven manner.

By comparing ATAC-seq profiles from T2D and ND donors, we identified 980 and 902 regulatory elements that exhibit quantitative T2D-associated increases or decreases in chromatin accessibility, respectively. These data suggest that T2D state by itself may not lead to widespread changes in chromatin accessibility. However, we acknowledge that T2D-associated epigenomic changes may be masked by multiple factors, including: 1) the small and genetically heterogeneous islet cohort analyzed; (2) cell type-specific changes that are hidden by other islet constituent cells; and (3) steady-state, normoglycemic culture conditions that may mask changes elicited by the diabetic milieu. Moving forward, studies that account for these potential confounders in larger, genetically-stratified islet cohorts will be necessary to further confirm these T2D-associated changes and identify novel ones.

## Methods

### Study subjects and primary islet culture

Fresh human cadaveric pancreatic islets were procured from ProdoLabs or the Integrated Islet Distribution Program (IIDP). Upon arrival, cells were transferred into PIM(S) media (ProdoLabs) supplemented with PIM(ABS) (ProdoLabs) and PIM(G) (ProdoLabs) and kept in a T-150 non-tissue culture treated flask (VWR) for recovery at 37 C and 5% CO_2_ overnight. ATAC-seq and RNA-seq were performed the following day as described below. For genotyping, genomic DNA was collected from islets cultured in CMRL + 10% FBS + Pen/Strep + Glutamax (Life Technologies) on tissue treated T175 until confluent and then prepped with the Qiagen Blood and tissue kit.

### Islet genotyping and imputation

Islets were genotyped using the Illumina Omni2.5Exome (n=11) or the Omni5Exome chips (n=8) (See Table S1). Genotype calls were made using the Genome Studio software (Illumina). The resulting vcf files were merged using the vcf-merge command in the vcftools/0.1.12a suite, and subsequently filtered for sites with any missing data (-- max-missing 1). 2.38 million genotyped SNVs passed QC and were used for imputation (1000G Phase 3 v5) (1000 Genomes Project Consortium et al. 2015) and phasing (Eagle v2.3) (Loh et al. 2016) using the Michigan Imputation Server (Das et al. 2016), to get a total of 47 million SNVs. After removing SNVs that were either monomorphic or outside islet ATAC-seq peaks, 1.21 million SNVs were kept for downstream analysis (caQTL/eQTL).

### Chromatin accessibility analysis (ATAC-seq)

Human islet ATAC-seq libraries were prepared as described (Varshney et al. 2017). Approximately 50-100 islet equivalents (50,000-100,000 cells) per sample were transposed in triplicate. Libraries were sequenced on an Illumina NextSeq500 (see Table S2). Paired-end 75-bp ATAC-seq reads were trimmed to remove low quality base calls using trimmomatic, and aligned to the hg19 human genome assembly with the Burrows Wheeler Aligner-MEM (Li and Durbin 2009). For each sample, duplicates were removed and the residual reads were shifted as previously described (Ucar et al. 2017). For each sample, technical replicates were merged using samtools, and peaks were called from the resulting merged bam file for each individual using MACS2 (Zhang et al. 2008) (with parameters -callpeak --nomodel -f BAMPE). Islets with less than 30,000 peak calls were removed, resulting in 19 islets for downstream analyses. An average sequencing depth of 62.6 million (SD=18.6 million) reads was obtained for each of the remaining 19 islets, after merging the 3 technical replicates. ATAC-seq peaks on sex chromosomes and those overlapping regions with low mappability(http://hgdownload.cse.ucsc.edu/goldenpath/hg19/encodeDCC/wgEncodeMapability/) were removed. The remaining autosomal peaks with q-values < 0.01 were selected for downstream analysis. The R Diffbind package (Stark and Brown) was used to define 154,437 ATAC-seq peaks for the 19 islets and to obtain read counts for each ATAC-seq peak for all the samples.

### Islet chromatin accessibility quantitative trait locus (caQTL) analyses

VerifyBamID (Jun et al. 2012) was used to match ATAC-seq bam files for each sample to each individual’s genotypes and ensure no samples were switched. We removed 69,939/154,438 islet ATAC-seq peaks containing monomorphic SNVs from the analyses. For the 1.21 million SNVs that were non-monomorphic and found within the remaining 84,499 islet peaks, allele-specific counts were obtained. Along with the read count information for the islet peaks for each sample local caQTLs were mapped using the RASQUAL statistical approach. The first 5 principal components were used as covariates to minimize confounding factors. The Bonferroni method was used to correct for the number of SNVs tested for each ATAC-seq peak. 10 random permutations were generated for each feature, and used to correct for the number of features tested, with an FDR cutoff of 10%.

### Differential ATAC-seq peak analyses (T2D vs. ND)

Islets from 5 T2D individuals and from 5 ND individuals with the best demographic match (e.g., age, sex and race) were selected for comparative/differential analysis (see Table 1). The R Diffbind package (Stark and Brown) was used to define 117599 consensus ATAC-seq peaks among these 10 islets and to determine read counts in each ATAC-seq peak for each of the 10 samples. Peaks were excluded from differential analysis if they met the following criteria: 1) the peak is present in fewer than three islet donors; 2) it is present in only 1 T2D and 2 ND (or 1ND and 2 T2D) islet donor. Surrogate Variable Analysis (SVA) (Leek et al. 2012) was used to summarize sources of unwanted variability in the read count table for the remaining 52,387 consensus peaks. The two significant surrogate variables were used as covariates in the design matrix to minimize confounding factors. The edgeR package (Robinson et al. 2010) was used to identify 1882 differential peaks at FDR 10%.

### Enrichment of genome-wide association study (GWAS) SNVs in differential (T2D vs. ND) and caQTL open chromatin sites

Lists of reference SNV identifiers were obtained from the NHGRI-EBI Catalog of GWAS SNVs (https://www.ebi.ac.uk/gwas/; accessed on January 19th, 2017) for 642 disease categories. For each disease category, GWAS SNVs were pruned using PLINK (Purcell et al. 2007) version 1.9 and parameters “--maf 0.05 --clump --clump-p1 0.0001 --clump- p2 0.01 --clump-r2 0.2 --clump-kb 1000” to ensure that each variant haplotype was tested only once during the enrichment analysis. For each SNV pair in linkage disequilibrium (LD) (R^2^ > 0.2) the SNV with the less significant p-value was discarded. GREGOR (Schmidt et al. 2015) was used to determine if the LD-pruned GWAS SNVs were enriched (r^2^ >0.8) in (1) differential or (2) caQTL ATAC-seq peaks. Diseases for which there weren’t any GWAS SNVs in LD (r^2^ >0.8) with the tested genomic regions were excluded from downstream analysis.

### Transcription factor (TF) motif enrichment

The findMotifsGenome.pl (with parameters hg19 and –size given) script in the Homer suite (Heinz et al. 2010) was used to identify significantly enriched transcription factor (TF) motifs in islet ATAC-seq data. For **Figure 1F**, motifs enriched in the islet-specific ATAC-seq peaks were identified using the common ATAC-seq peaks as the background set. Common (background) peaks were defined as those that overlapped any given ATAC-seq peaks for adipose, CD4T, GM12878 and 2 PBMC samples. Islet-specific peaks were those that did not overlap an open chromatin loci in the other tissues. For **Figure 4G**, motifs enriched in the 1882 differential peaks were identified using the 52387 peaks as the background set. For **Figure 2E**, motifs enriched in the 3001 caQTL peaks were identified using the 154437 peaks as the background set. TFs are clustered based on the similarity of their Position Weight Matrices (PWMs) using Kullback Leibler divergence method as implemented in the TFBSTools R package (Tan and Lenhard 2016).

### ChromHMM annotation

Harmonized ChromHMM files (13 state) for islets, the ENCODE cell lines and the Roadmap tissues were used as previously determined (Varshney et al. 2017). The ggplot2 package was used to plot the overlap of peak sets to the harmonized ChromHMM states. For cases when a peak overlapped two or more ChromHMM states, the order of preference for overlaps were as follows: Active TSS, Bivalent TSS, Weak TSS, Flanking TSS, Active Enhancer-1, Active Enhancer-2, Weak Enhancer, Genic Enhancer, Strong Transcription, Weak Transcription, Repressed Polycomb, Weak Repressed Polycomb, and Quiescent.

### Stretch enhancer annotation

Stretch enhancers were defined using the harmonized ChromHMM definitions. Briefly, stretch enhancers are defined as > 3kb consecutive segments that overlap and enhancer state including Active Enhancer 1 and 2, Weak Enhancer and Genic Enhancer ChromHMM states. To test whether a peak set is enriched in a given tissue stretch enhancers, fisher’s exact test was performed. The background set was the union of the stretch enhancers of all 31 tissues, except the one being tested.

### RNA-seq profiling

Libraries for the 19 islets exhibiting high-quality ATAC-seq profiles were prepared using the stranded TruSeq kit (Illumina), and had either ERCC Mix 1 or Mix 2 randomly spiked-in (ThermoFisher, catalog #4456740; see Supplemental Table S3). The 10 islets used for the T2D vs. ND differential analysis were sequenced on an Illumina NextSeq500 sequencer. The remaining 9 islets were sequenced separately on Illumina HiSeq 2500. The paired-end RNA-seq reads for each islet was trimmed for low quality base calls using trimmomatic (Bolger et al. 2014). Bowtie2 (Langmead and Salzberg 2012), in conjunction with RSEM (Li and Dewey 2011) (rsem-calculate-expression), was used to obtain the FPKM and Expected read counts for all genes across the 19 samples. An average depth of 87.2 ± 27.8 million reads was obtained for the 19 islets.

### Differential gene expression

Only autosomal genes with FPKM>5 in more than 3 islets were included in the analysis. SVA was used to summarize sources of unwanted variability in the expected read count matrix for the remaining 10,116 genes, and minimize confounding factors. Differential gene expression analyses between ND and T2D samples were completed using edgeR at FDR 10%.

### Expression QTLs analysis

Expected counts from RSEM for 9656 genes expressed (FPKM>5) in the 19 islets were used as input to RASQUAL. Only SNVs within the genes or those flanking 50 kilobases (kb) on either side of the gene body were tested for eQTL activity. The first four principal components and race were used as covariates to minimize confounding factors. Bonferroni correction was used to correct for the number of SNVs tested for each gene. 10 random permutations was generated for each gene, and used to correct for the number of genes tested, with an FDR cutoff of 10%.

### Luciferase reporter assays

Genomic DNA from individuals homozygous for the reference and alternate alleles was used to amplify the 9 loci (see Supplemental Table S4). The 18 total constructs were cloned into the pDONR vector with BP Clonase (Invitrogen), which was then used to transfer the constructs into the Gateway-modified pGL4.23-FWD vector (Stitzel et al. 2010) with LR Clonase. Renilla luciferase (pRL-TK) was co-transfected with equimolar amounts of each pGL4.23 vector into MIN6 was used to normalize differences in transfection efficiencies as previously described (Stitzel et al. 2010). Cells were lysed in 1x Passive Lysis Buffer (PLB) 36 hours after transfection and luciferase activity was measured using the Dual Luciferase Reporter (DLR) Assay system (Promega) according to the manufacturer’s instructions. DLR activity was measured using a Synergy2 Microplate Reader (BioTek). The DLR ratio (Firefly/Renilla) for each construct was normalized to the empty pGL4.23 vector. The assay was performed 3 times. Each run included 3 separate mini-preps for each construct, and 3 technical replicates for each mini-prep.

## Supplementary Figure Legends

**Figure S1. Chromatin accessibility maps in human islets. (A)** Insert size distributions of six representative islets (3 non-diabetic (ND), 3 T2D). ATAC-seq libraries capture nucleosome free and mono-, di-nucleosomal regions. (**B**) Number of ATAC-seq peaks called across the cohort, ranging from individual-specific peaks (n=1) to common peaks (n=19). (**C**) Islet ChromHMM annotations for ATAC-seq peaks categorized with respect to their frequency in the cohort. Note that common peaks are mostly promoters, whereas individual-specific or rare peaks include more quiescent or low signal regions. (**D**) TF motifs enriched in islet-specific peaks (from Figure 1F).

**Figure S2. Chromatin accessibility QTLs in islets. (A)** QQ-plot for expected and observed caQTL p-values. (**B**) Location of caQTLs (marked in green) across the genome. Note that caQTLs are widely distributed across each chromosome. (**C**) Functional annotation of caQTLs using ChromHMM states in islets and other tissues. Tissues are sorted from highest to lowest overlap between ATAC-seq peaks and the ‘Quiescent/Low Signal’ state. Note the enrichment of caQTLs in islet enhancers. (**D**) Overlap of caQTLs with islet eQTLs from the same cohort. (**E**) Overlap of caQTLs with previously published islet eQTLs from 112 individuals (Varshney et al. 2017). (**F**) Table enumerating the 9 caQTLs that were tested for luciferase activity. Note that higher accessibility is associated with higher enhancer activity for all the 5 caQTLs displaying differential luciferase activity.

**Figure S3. T2D-associated chromatin accessibility changes. (A)** Principal Components 1 and 2 for the 10 islets. Note that the ND and T2D islets do not cluster together using all ATAC-seq peaks. (**B**) The weighted average proportion variance explained for the meta-variables associated with the 10 islets. Note that SVA reduces the variance attributed to all meta-variables, except for the one of interest (Condition) (**C**) The overlap of differential peaks detected with and without SVA is significant. (**D**) MA plot of all ATAC-seq peaks used for differential analyses (n=52,387). Positive logFC means the peaks are opening in T2D, and Negative logFC means the peaks are closing in T2D (CPM=counts per million). (**E**) Each cell in the heat map shows the number of differential peaks that are called as ATAC-seq peaks among ND and T2D islets. 918/1882 differential peaks are found in all 10 islets. (**F**) Venn diagram showing the number of opening or closing peaks overlapping caQTLs.

## Supplementary Table Legends

**Table S1.** Meta data associated with the 19 islets.

**Table S2.** ATAC-seq quality control metrics for the 19 islets.

**Table S3.** RNA-seq quality control metrics for the 19 islets.

**Table S4.** Constructs for Luciferase Assay

**Table S5.** Differentially accessible ATAC-seq peaks in Islets

## Acknowledgements

We are indebted to the anonymous pancreatic organ donors and their families, without whom this study would not be possible. We thank Jane Cha for her help in generating artwork for the figures and members of the Stitzel and Ucar labs for helpful discussion and critiques during study design, completion, and write-up. This study was supported in part by the National Institute of Diabetes and Digestive and Kidney Diseases (NIDDK) under award number DK092251 (to MLS) and by The Jackson Laboratory Startup and Director’s Innovation Funds (to M.L.S. and D.U.). Opinions, interpretations, conclusions, and recommendations are solely the responsibility of the authors and do not necessarily represent the official views of the National Institutes of Health.

